# Culture Associated DNA Methylation Changes Impact on Cellular Function of Human Intestinal Organoids

**DOI:** 10.1101/2022.04.25.489354

**Authors:** Rachel D Edgar, Francesca Perrone, April R Foster, Felicity Payne, Sophia Lewis, Komal M Nayak, Judith Kraiczy, Aurélie Cenier, Franco Torrente, Camilla Salvestrini, Robert Heuschkel, Kai O Hensel, Rebecca Harris, D. Leanne Jones, Daniel R Zerbino, Matthias Zilbauer

## Abstract

**Background & Aims:** Human intestinal epithelial organoids (IEO) are a powerful tool to model major aspects of intestinal development, health and diseases, as patient derived cultures retain many features found *in-vivo*. A necessary aspect of the organoid model is the requirement to expand cultures *in-vitro* through several rounds of passaging. This is of concern, as the passaging of cells has been shown to affect cell morphology, ploidy, and function. In this study, we address concerns around long term passaging of IEO to better characterise and define effects on cell morphology and function.

**Methods:** Here we have analysed 173 human IEO from two sampling sites, terminal ileum and sigmoid colon and examined the effect of culture duration on DNA methylation (DNAm), gene expression and cellular function including their response to proinflammatory cytokines and *in-vitro* cell differentiation.

**Results:** Our analyses revealed a major effect of culture duration on DNAm, leading to significant changes at 61,337 loci representing approximately 8% of all CpGs tested. Although global cellular functions such as gut segment-specific gene expression remained stable, a subset of methylation changes correlated with altered gene expression at baseline as well as in response to inflammatory cytokine exposure and *in-vitro* differentiation. Importantly, epigenetic changes were found to be enriched in genomic regions associated with colonic cancer and distant to the site of replication indicating similarities to malignant transformation.

**Conclusions:** Our study reveals culture-associated epigenetic, transcriptomic and functional changes in human mucosa derived IEO and highlights the importance of considering passage number as a potentially confounding factor.

**Synopsis:** This work describes cell culture induced changes to DNA methylation, gene expression and cellular function in human IEO. Globally organoids lost DNA methylation with time in culture while DNA methylation also became generally more variable. This work suggests a shifted epigenetic profile in organoids cultured long-term.

## INTRODUCTION

Organoids are self-organising three-dimensional structures that are derived from either pluripotent or somatic, organ specific, stem cells. They have been shown to closely mimic both anatomy and cellular function of the *in-vivo* organ. Importantly, the ability to generate such organoids from human cells has turned them into powerful translational research tools with a wide range of applications, including the development of new therapeutics, testing of existing drugs as well as the application of a personalised treatment approach in several diseases^1–5^.

Amongst the most advanced human organoid models, are mucosa derived intestinal epithelial organoids (IEO)^6^. Isolation of LGR5+ intestinal stem cells or entire crypts followed by their culture in an environment closely mimicking the *in-vivo* stem cell niche leads to the development of three dimensional mini-organs containing all epithelial cell subsets organised in a crypt-villus structure, closely reflecting the *in-vivo* situation. Mucosal IEO have been successfully generated from all parts of the digestive tract and from a wide range of donors including healthy individuals of different age groups and patients with intestinal diseases such as Inflammatory Bowel Disease (IBD)^7–11^. The latter provides unprecedented opportunities to investigate disease pathogenesis and develop novel treatment approaches using patient derived organoids.

Another major advantage of human IEO is the ability to keep them in culture over prolonged time periods (i.e. months or even years). Indeed, prolonged culture periods are also required to sufficiently expand organoids prior to their experimental use. As a result, cellular cultures must undergo numerous rounds of ‘passaging’, a process during which individual organoids are dissociated into individual crypts that then give rise to new organoids, thereby increasing their total number. The potential impact of prolonged *in-vitro* culturing of a human mucosa derived IEO remain largely unknown. Although previous studies have demonstrated a high degree of genetic stability over time^12^, little is known about epigenetic alterations or changes in cellular function. Despite major progress in optimising culturing methods aimed at closely mimicking the *in-vivo* situation, fundamental differences such as the absence of gut microbiota or signalling from other cell types (e.g. immune or mesenchymal cells) may cause alterations in both cellular epigenome and/or function, as has been shown in other cell culture models^13,14^. Understanding the potential impact of prolonged culturing and repeated passaging on cellular function of human IEO is, therefore, of critical importance as any changes may confound experimental results as well as impact on their potential use in the field of regenerative medicine.

DNA methylation (DNAm) is one of the main epigenetic mechanisms known to play a key role in regulating cellular function of human cells including the intestinal epithelium^7,8,15–18^. We have previously reported that gut segment specific DNAm signatures are faithfully retained in human mucosa derived IEO and that they are critical for region specific cellular function^10,19^. Although such gut segment specific DNAm signatures were found to be highly stable over time in IEO derived from children and adults, human fetal gut derived IEO displayed substantial changes in their DNAm profiles during prolonged culturing suggesting a degree of *in-vitro* maturation.

Importantly, epigenetic instability in intestinal cells is also seen in colon carcinogenesis, with substantial genome-wide DNAm changes observed in colorectal cancer (CRC)^20^. Specifically, when compared to healthy colonic mucosa, CRC shows both losses and gains of DNAm as well as increased variability in DNAm^20–22^.

Here we set out to monitor global DNAm in human IEO during prolonged *in-vitro* culture and investigate the impact of associated epigenetic changes on cellular function. Based on the analyses of 173 human, mucosa derived IEO, we have identified distinct culture associated DNAm changes, some of which impact gene transcription and cellular function. This highlights the importance of considering culture duration in experimental design and interpretation of results.

## RESULTS

### Prolonged *in-vitro* Culture of Human IEO is Associated with Distinct DNAm Changes

In order to examine the potential impact of prolonged *in-vitro* culture on human IEO DNAm, we recruited a total of 46 children undergoing routine endoscopy and obtained mucosal biopsies from the distal small bowel (i.e. Terminal Ileum (TI)) and distal large bowel (Sigmoid Colon (SC)). Human IEO were generated (n=80, Cohort 1, **Table 1**) and cultured over several months whilst documenting the number of passages as an indication for culture duration (**Figure 1A**). IEO were harvested at various time points ranging from passage 1 (approximately 7-10 days) to passage 16 (approximately 4-10 months culture duration), and genome wide DNAm profiling was performed. First, we performed principal component analysis of IEO DNAm profiles. As shown in **Figure 1B**, IEO DNAm was strongly associated with gut segment (also **Figure 1C**), age and gender as expected. Interestingly, culture duration (measured by passage number) was found to be significantly associated with variation in PC1- 3 and 6 (**Figure 1B and E**). As reported previously, DNAm of IEO is highly gut segment specific with differences between gut segments (i.e. small bowel versus large bowel) being preserved even over prolonged culture duration (**Figure 1C and D**). Only 4.4% CpGs with significant segment specific DNAm (29,805 CpGs; FDR<0.0001, a difference in mean DNAm between segments or |delta beta| >0.2) were also seen to change with passage (**Figure 1D**). Similarly, biological age of the donor was retained, as it was accurately reflected when calculating epigenetic age^23^ of IEO (r_s_= 0.52; **Figure S1**), even in high passage IEO, suggesting epigenetic age is maintained across passaging.

**Table 1.** Intestinal Organoid datasets used in analysis. Cohorts are either generated for this analysis or are publicly available from the listed sources.

| Cohort | Sample |  | Gut |  | Age | Gender |  |
| --- | --- | --- | --- | --- | --- | --- | --- |
|  | Number | Data Type | Segments | Passage |  | (%F) | Source |
| 1 | 80 | DNAm | TI and SC | 1-16 | Pediatric | 55 | Newly generated |
| 2 | 30 | DNAm | TI and SC | 1-11 | Pediatric | 50 | E-MTAB-4957 [19] |
| 3 | 21 | DNAm | Colon,<br>Duodenum,<br>Jejunum | 2-11 | Pediatric<br>and Adult | 48 | GSE141256 [10] |
| 4 | 42 | DNAm<br>and gene<br>expression | TI and SC | 2-12 | Pediatric | 67 | Newly generated |

**Figure 1:**
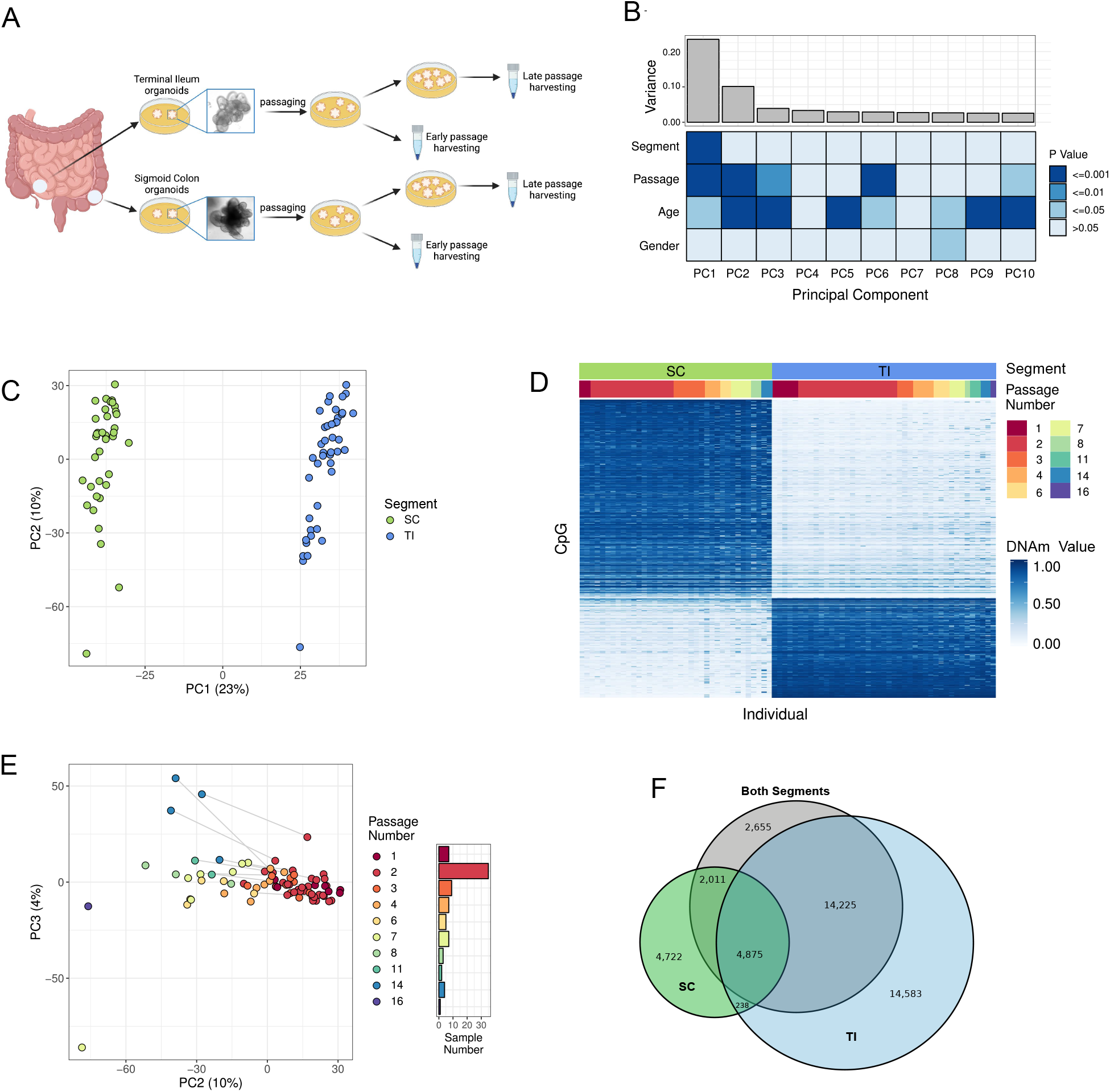
Sampling site of origin and IEO passage number are associated with the main components of DNAm variation. (A) Outline of study design. (B) The scree plot shows the amount of DNAm variance accounted for by each PC. The heat map shows the associations between sample variables and each PC. P values were generated with a Spearman correlation for continuous variables or an ANOVA for categorical. (C) PC1 and PC2 are plotted for each sample. Samples are coloured by the segment of origin. (D) DNAm of the top 500 CpGs, in rows, differentially DNAm between gut segments. IEO are sorted, in columns, by segment then by passage. (E) PC2 and PC3 are plotted for each sample. Samples are coloured by passage number. Lines connect samples derived from the same patient, but where IEO were cultured to a different number of passages. The histogram in the legend shows the distribution of passage numbers across the cohort. (F) Overlap of CpGs differentially DNAm in the combined cohort of both segments, and in the TI or SC separately.

However, when considering the entire DNA methylome, 10% of variability (i.e PC2) was found to be strongly associated with the number of passages (r_s_= -0.82**; Figure 1E**). When performing analyses in IEO divided by gut segment of origin, small bowel (TI) and large bowel (SC), we observed a major overlap in the CpGs that changed their DNAm pattern over time (**Figure 1F**), suggesting approximately half of the passage associated epigenetic changes are independent of gut segment.

Together these results demonstrate that whilst the vast majority of DNAm appears to be stable even during prolonged culture periods, approximately 8% of CpGs (61,337 CpGs) display significant culture associated DNAm changes, independent of gut segment.

### Passage Associated DNAm Changes Validate in Additional IEO Culture Cohorts

In order to ensure that observed culture associated DNAm changes were not a result of laboratory or sample cohort specific culturing techniques, we tested this phenomenon in additional, publicly available cohorts generated by our group (Cohort 2) as well as by an independent group (Cohort 3, **Table 1**). In both additional cohorts, IEO were generated from mucosal biopsies obtained from the small (Terminal Ileum, Duodenum, Jejunum) and large bowel (Colon) and passage number recorded. Genome wide DNAm was assessed using Illumina arrays (450K and EPIC). Performing the same principal component analysis on Cohorts 2 and 3 confirmed highly significant DNAm changes associated with passage (**Figure 2A and C**). Looking at individual CpGs, out of 23,766 displaying passage-associated differential DNAm in the discovery Cohort 1, approximately 7,600 CpGs were present on the arrays used in Cohorts 2 and 3. Strikingly, 4,748 (62%) and 4,620 (60%) of Cohort 1 passage associated CpGs were also found to change DNAm with passage number in Cohorts 2 and 3 respectively. Moreover, direction of change (i.e. DNA hypo/hyper-methylation) associated with passage were highly consistent showing a major overlap in both validation cohorts (**Figure 2B and D**).

**Figure 2:**
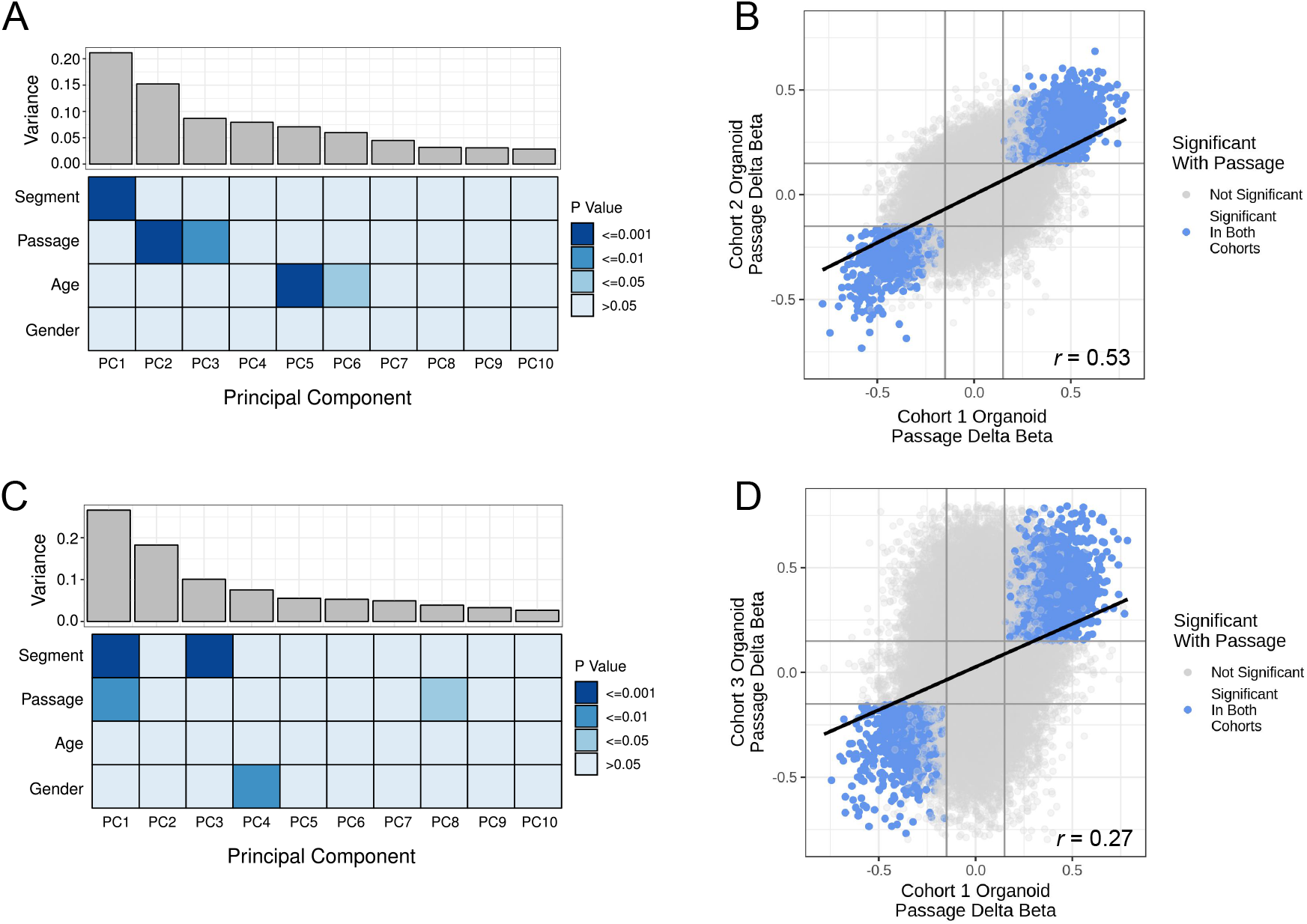
Long term culture effects on IEO DNAm are validated in independent cohorts. (A) and (B) Cohort 2 (C) and (D) Cohort 3. (A) and (C) The scree plots show the amount of DNAm variance accounted for by each PC. The heat map shows the associations between sample variables and each PC. P values were generated with a Spearman correlation for continuous variables or an ANOVA for categorical. (B) and (D) Direction of effect is consistent between cohorts. The delta betas from two cohorts are shown as points with CpGs significantly associated with passage highlighted.

Taken together, passage associated DNAm changes were validated in two additional cohorts further confirming that this phenomenon occurs independent of cohort and laboratory.

### High and Low Passage IEO Display Distinct Transcriptional and Functional Differences, Some of Which are Associated with DNAm Changes

Having observed major DNAm changes associated with prolonged culturing of IEO, we next aimed to investigate the potential impact of culture duration on cellular function and gene transcription. We therefore generated an additional cohort (Cohort 4) of IEO from small and large bowel biopsies from 5 healthy individuals (**Figure 3A**). IEO were kept in culture for up to 12 passages (approximately 4 months) and subjected to a range of functional assays as well as genome wide epigenetic and transcriptomic profiling. As shown in **Figure 3B**, culture duration did not impact on IEO appearance and distinct gut segment specific threedimensional differences were retained with small bowel IEO demonstrating a more budded appearance whilst colonic IEO appear overall more cystic (**Figure 3B**). Similarly, measuring IEO size over several days after passaging demonstrated no difference in growth when comparing IEO at late versus high passage (P<0.05; **Figure 3C)**.

**Figure 3:**
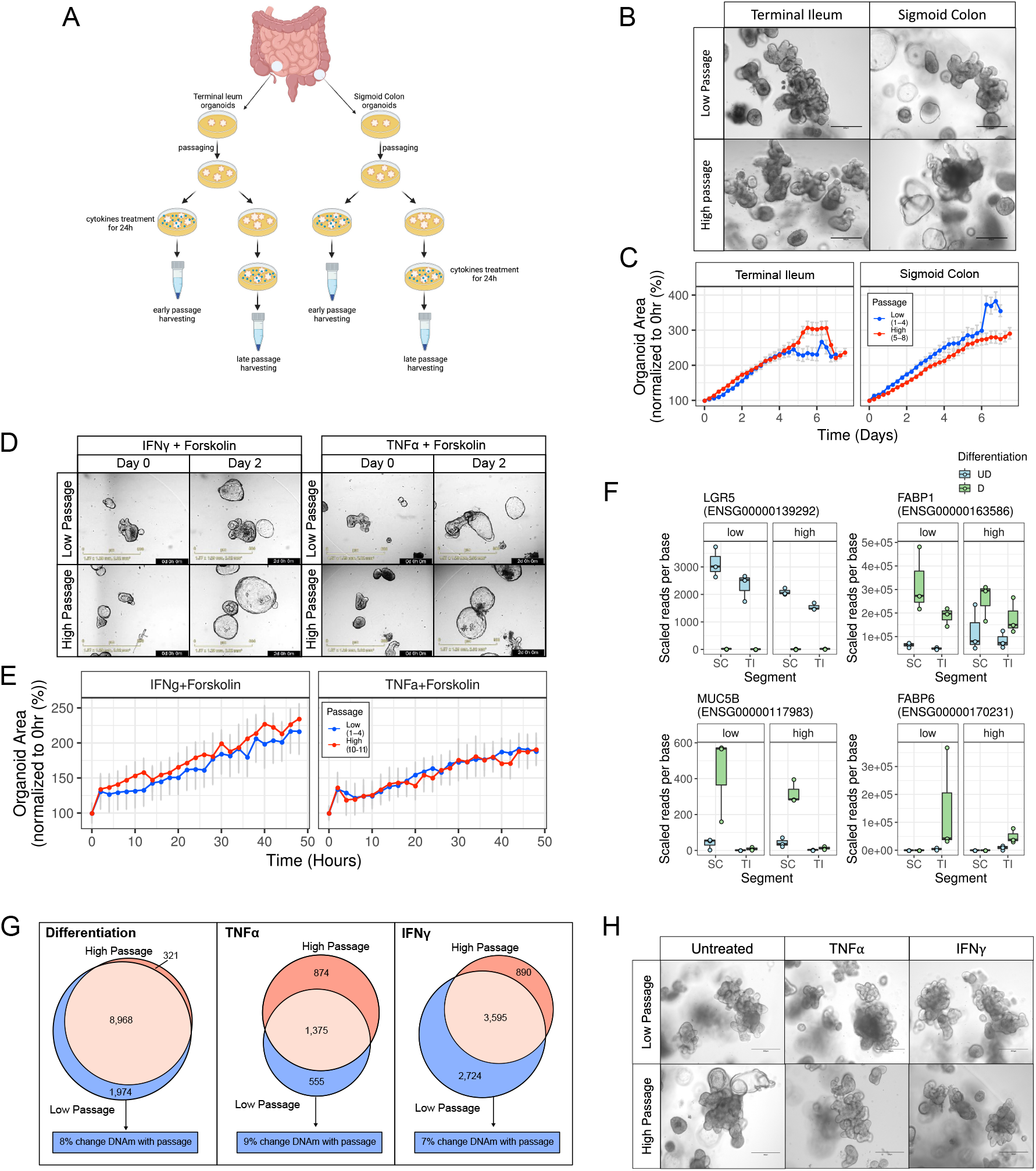
High and low passage IEO are functionally similar. (A) Outline of IEO function experimental design. (B) Bright-field images of TI and SC derived IEO at low and high passage taken by EVOS FL system (Life Technologies), scale bars= 300*μ*m. (C) Comparison of the growth curves between low and high passaged TI and SC derived IEO. The curves were generated following the analysis of IEO area with Incucyte IEO analysis software, from images taken every 6 hours over 7 days for each passage (n=2, 6 technical replicates per each biological replicate).(D) Representative images of TI organoids treated with IFN*γ*+Forskolin and TNF*α*+Forskolin taken with by Incucyte, scale bars= 800*μ*m. (E) Comparison of the organoids area between early and late passage organoids following pro-inflammatory cytokines and Forskolin treatments (n=2, 4 technical replicates per each biological replicate. (F) Representative differentiation marker genes. Gene expression is shown for each gene split by gut segment and passage (high or low) of the IEOs. Boxplots are colored by the IEO differentiation status. (G) Overlap, between high and low passage IEO, of genes significantly differentially expressed upon differentiation, IFN*γ* and TNF*α* stimulation. For the genes only differential in low passage IEO the proportion of these also differentially DNAm with passage is given. (H) Bright-field images of TI derived IEO at low and high passage following proinflammatory cytokines treatment (TNF*α* or IFN*γ*), taken by EVOS FL system (Life Technologies), scale bars= 300*μ*m.

Next, we aimed at testing whether culture duration impacts barrier function of IEO at baseline and in response to inflammatory cytokines. We therefore developed a modified Forskolin-Induced Swelling assay, which has been used previously in the context of cystic fibrosis in order to test the function of cystic fibrosis transmembrane conductance regulator (CFTR) in gut organoids^24,25^. Briefly, the assay takes advantage of the ability of Forskolin to increase cyclic adenosine monophosphate (cAMP) levels in the intestinal epithelium. This in turn results in the opening of iron channels followed by transport of ions and water into the lumen of organoids causing them to swell up. IEO with intact barrier will swell up over time whilst any damage to barrier function will either stop IEO from swelling and or limit their capacity of swelling. IEO size therefore correlates with barrier function (**Supplementary Video 1**). As shown in Supplement Figure S3, IEO incubated with either TNFα or IFNγ and Forskolin for 48h show significantly reduced swelling compared to IEO incubated with Forskolin only (P<0.05), suggesting that these cytokines impact on epithelial barrier function as expected (Figure S3). Importantly, culture duration did not impact on epithelial cell barrier as we did not observe any difference between high and low passage IEO (P>0.05; **Figures 3D and E**).

In order to test the potential impact of culture duration on the ability of the human intestinal epithelium to differentiate into cell subsets, IEO were subjected to *in-vitro* differentiation by withdrawing Wnt agonists over 4 days^19^. Gene transcription was assessed on extracted RNA using RNA sequencing. As described previously, *in-vitro* differentiation led to decreased expression of intestinal epithelial stem cell marker *LGR5* whilst expression of epithelial cell subset (*MUC5B*) and differentiation markers (*FABP1* and *FABP6*) increased in a gut segment specific manner (**Figure 3F**). Importantly, passage number did not impact the expression of selected gut segment specific marker genes, or on the microscopic appearance of IEO (**Figure S4**). Furthermore, out of 10,942 genes found to change expression in response to *in-vitro* differentiation in low passage IEO, expression of 8,968 also changed in high passage IEO (FDR < 0.05). Only 8% of the genes which change during differentiation in low but not high passage IEO, are associated with at least 1 CpG demonstrating passage associated DNAm changes (**Figure 3G left panel; Table S7**). At a more stringent cut off for differential expression (FDR < 0.01 and |log fold change| > 0.5) 5,224 genes changed expression with differentiation in low passage IEO, and 2,790 of these also changed in high passage IEO. To evaluate the potential impact of culture duration on the responsiveness of IEO to inflammatory stimuli, low and high passage IEO were co-cultured with IFNγ or TNFα for 24 hours, and gene expression was assessed on extracted RNA (**Figure 3A**). As shown in **Figure 3G**, a major overlap was found between differentially expressed genes in response to IFNγ and TNFα comparing low with high passage IEO and no microscopical changes observed (**Figure 3H**). Similar to transcriptional changes in response to *in-vitro* differentiation, of the genes only changed in low passage IEO, 7 and 9% (IFNγ and TNFα respectively), were associated with at least 1 CpG displaying passage associated DNAm changes, indicating a limited impact of culture associated epigenetic changes on inflammation induced gene transcription. However, a highly significant association between passage associated DNAm and transcriptional changes was observed for several genes including *EDAR* and *EIF4G1* (**Figure 4A and S5; Table S6)**. In addition, some of the genes where DNAm associated with passage show striking differences in gene expression in response to differentiation and stimulation (**Figure 5B; Table S8 and S9**). Interestingly, transcriptional responses to both *invitro* differentiation and exposure to IFNγ were larger in early versus late passage IEO suggesting culture duration may impact on the overall magnitude of response in the intestinal epithelium.

**Figure 4:**
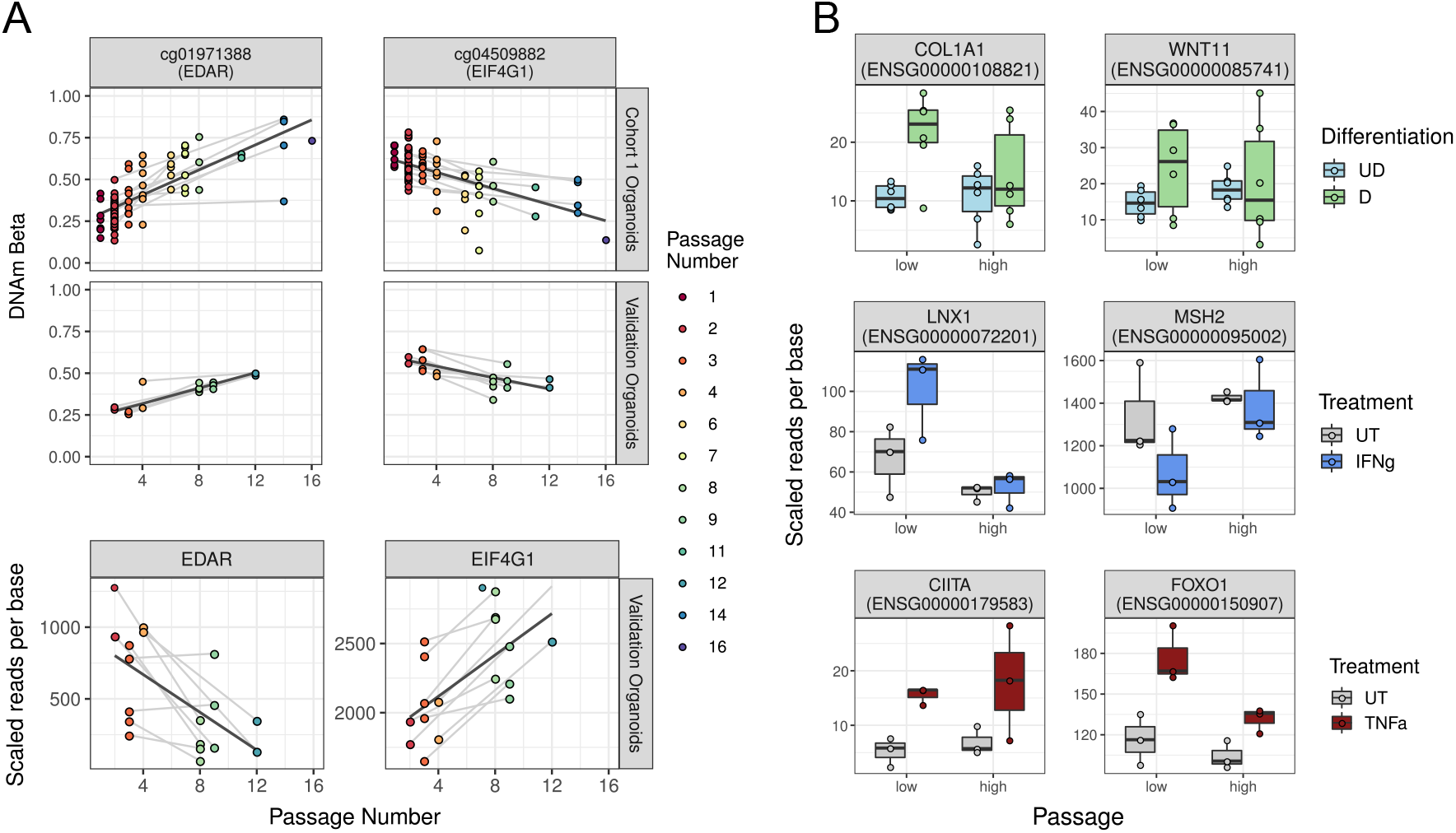
Genes are differentially expressed with IEO passage. (A) Representative CpGs and genes with DNAm and expression significantly associated with passage in Cohort 1 and the validation cohort. Samples are coloured by passage number and grey lines connect samples derived from the same patient. Regression lines between passage and DNAm/expression are in black. (B) Representative genes with a difference in response to differentiation, IFN*γ* or TNF*α* between low and high passage IEOs.

**Figure 5:**
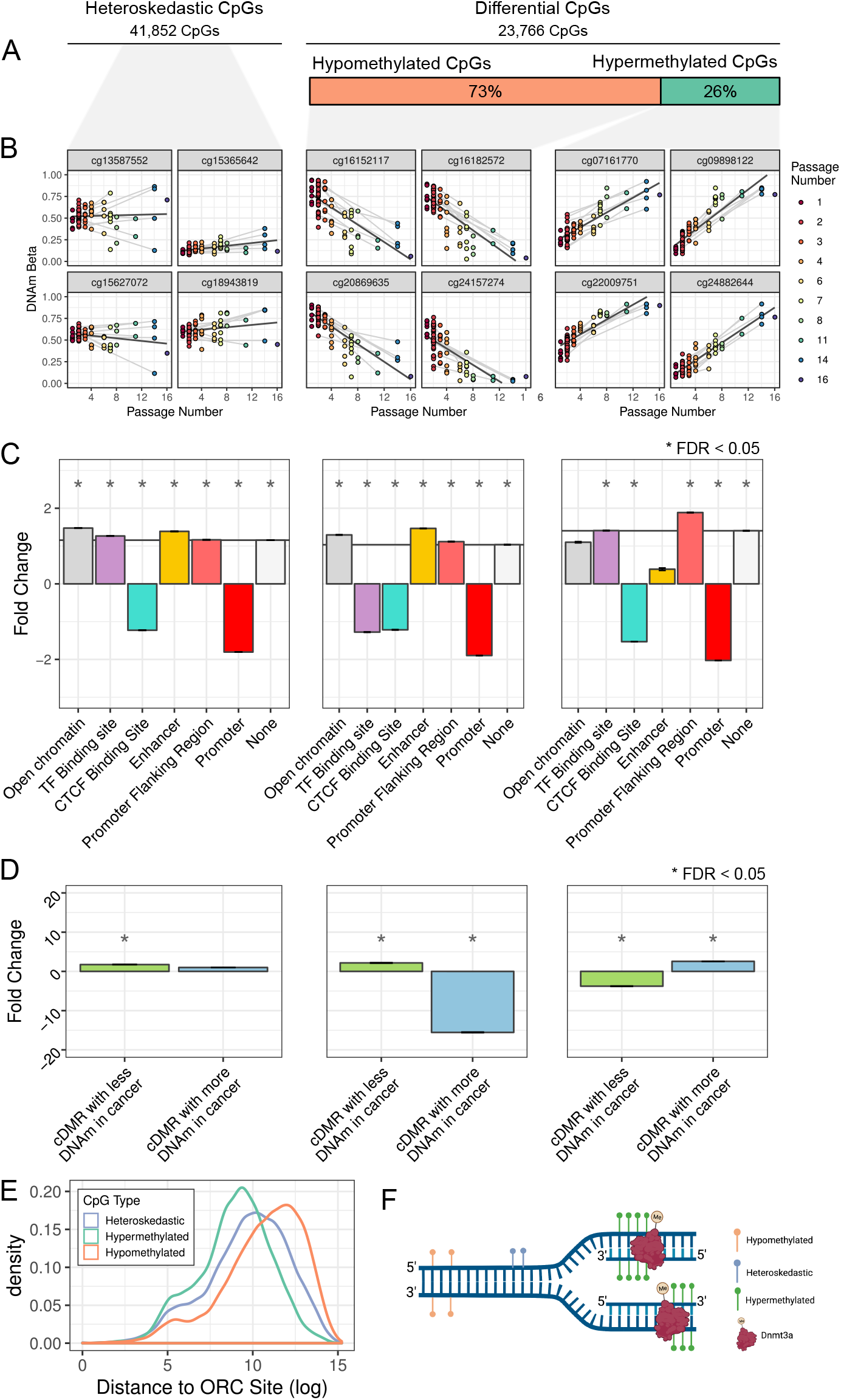
Passage affects DNAm in specific regions of the genome. (A) Proportion of passage CpG split by type of change in DNAm. (B) Representative CpGs with DNAm significantly associated with passage. CpGs were selected as either significantly heteroskedastic with passage or are differentially DNAm, either losing (hypomethylated) or gaining (hypermethylated) DNAm with increasing passage. Samples are coloured by passage number and grey lines connect samples derived from the same patient. Regression lines between passage and DNAm are in black. (C) Fold change between the number of passages associated CpGs in regulatory genomic features and expected number based on the EPIC array background. Standard error bars around the mean fold change are for the error across 1,000 random samplings. (D) Fold change between the number of passages associated CpGs in cDMRs and expected number based on the EPIC array background. Standard error bars around the mean fold change are for the error between 1,000 random samplings. (E) Distance of passage associated CpGs from ORC sites. (F) Schematic of passage DNAm changes with origins of replication.

In summary, although the impact of culture duration on microscopic appearance and gross cellular function of IEO appears to be limited, distinct transcriptional changes were observed and some of them are associated with DNAm.

### Prolonged Culture of IEO Primarily Causes Global Loss of DNAm and Excludes Hypomethylated Promoter Regions

Having determined the impact of passage associated DNAm changes on gene transcription and cellular function of IEO, we next aimed to investigate specific patterns of epigenetic changes and their distribution across the genome.

In total we identified 61,337 CpGs with DNAm changes associated with IEO passage. Of these 23,766 CpGs (39%) showed significant differential DNAm with passage (FDR <0.05, |delta beta| > 0.15, **Figure 5A**). The majority of differential DNAm (i.e. 17,352, 73%) was a loss of DNAm whilst only 26% of CpGs (i.e. 6,414) gained DNAm during prolonged in-vitro culture (**Figure 5B**). A third category of culture associated DNAm changes at an individual CpG level are those that display a mixture of gain and loss of DNAm. This type of change, also referred to as heteroskedastic, has been described as the hallmark of epigenetic drift^26,27^. Epigenetic drift can be defined as the divergence of the epigenome as a result of age, caused by stochastic changes in DNAm. Interestingly, a total of 41,852 CpGs (5% of all CpGs measured and 68% of CpGs showing culture associated DNAm changes) were found to follow this pattern, suggesting a degree of epigenetic drift contributing to the observed passage associated DNAm changes (**Figure 5A and B**). Looking at the distribution of DNAm changes, heteroskedastic CpGs were enriched in transcription factor (TF) binding sites, open chromatin, enhancers and promoter flanking regions, but depleted in promoters and CTCF binding sites using established genome annotations^28^ (FDR <0.05, **Figure 5C**). Furthermore, CpGs losing DNAm were significantly enriched in open chromatin, enhancers and promoter flanking regions but depleted in promoters, CTCF binding sites and TF binding sites (FDR <0.05, **Figure. 5C**). In contrast, CpGs gaining DNAm in culture were enriched in TF binding sites and promoter flanking regions but also depleted in promoters and CTCF binding sites (FDR <0.05, **Figure. 5C**), suggesting that the dysregulation of DNAm with passage occurred genome wide, but seemed to spare non-variable unmethylated regions.

### Culture Associated DNAm Changes of Human IEO Share Features of Intestinal Cancer and Occur at Late Replicating Regions of the Genome

Global loss and local gain of DNAm has been linked to various malignancies including colon, breast and prostate cancer^29–32^. We next looked at a previously characterized list of differentially methylated regions previously seen in colon cancer (cDMRs)^20^. Interestingly, we found that within these cDMRs, passage associated DNAm changes in IEO followed the same pattern (i.e. gain or loss of DNAm) as those reported in colon cancer (**Figure 5D**). Moreover, CpGs that lost or gained DNAm in high passage IEO, were enriched in cDMRs that lose or gain DNAm in cancer, respectively (**FDR <0.05, Figure 5D**). Heteroskedastic CpGs were also enriched in cDMRs hypomethylated in cancer (**FDR <0.05, Figure 5D**).

It has been proposed in the context of cancer that late-replicating regions of the genome will have less time to remethylate CpGs on the daughter strand after DNA replication (**Figure 5E**)^33^. A replication associated loss of DNAm has also been observed in cell culture models^34^. We, therefore, tested if observed passage associated DNAm changes could be caused by inferred DNA replication timing. Using 52,251 origins of replication locations based on origin replication complex (ORC2) binding sites^35^ we found that CpGs that lose DNAm with passage were located further from origins of replication than expected by chance (P <0.001; **Figure 5F**). Interestingly, the same was true for heteroskedastic CpGs whilst hypermethylated CpGs were located closer to origins of replication than expected by chance (P <0.001; **Figure 5F**). Taken together these results suggest that passage-associated DNAm changes in human IEO share features of epigenetic changes observed in colon cancer and may at least in part be caused by fast cell turnover *in-vitro*.

## DISCUSSION

Since the establishment of human mucosa derived IEO as powerful translational research tools just over a decade ago, the number of applications has continued to increase dramatically. Major new areas of interest include precision and regenerative medicine, drug discovery and development, as well as modelling of disease pathogenesis^1–5^. Importantly, the vast majority of applications require IEO to be cultured for prolonged time periods in order to sufficiently expand cell numbers for larger scale experiments. Moreover, the ability to culture IEO over longer time periods (i.e. months) is also essential for the testing of specific chronic stimuli (e.g. inflammation or infection) on epithelial cell function over time. Although numerous studies have confirmed the ability to maintain human mucosa derived IEO in culture over prolonged time periods, there is still little information available on the potential impact of culture duration on cellular function.

As one of the main epigenetic mechanisms in mammalian cells, DNAm is known to play a key role in regulating intestinal epithelial cellular function. In this study we reveal major, culture associated DNAm changes in human, mucosa derived IEO occurring over several months, regardless from which gut segment or human donor the IEO were derived. Indeed, we were able to validate culture associated DNA methylation changes across several IEO cohorts that were generated both by independent groups, strongly suggesting this phenomenon is likely to apply to most, if not all mucosa derived IEO that are cultured using a similar, standard protocol as widely published^6,36^.

Given major differences in the microenvironment between the intestinal stem cell niche *in-vivo* and *in-vitro*, it is not surprising that we observe such epigenetic changes. However, their potential impact on cellular function is of critical importance to any downstream application.

Reassuringly, we found that the overall impact of culture associated DNA methylation changes in human IEO was limited and did not impact cellular identity (e.g. gut segment specific methylation signatures), epigenetic age or broad cellular function such as growth and intestinal barrier function. However, genome wide transcriptional analyses of IEO revealed distinct passage associated changes at baseline, as well as in response to *in-vitro* differentiation and exposure to inflammatory cytokines. Importantly, a subset of these were found to overlap with passage associated DNA methylation changes, strongly suggesting that epigenetic alterations impact on cellular function.

Our findings are in keeping with previous reports on directed and stochastic changes to DNAm with passaging of induced pluripotent stem cells (iPSCs)^37^. Specifically, iPSCs were reported to lose DNAm with increased passaging; tissue specific DNAm patterns observed in early passages were lost, and cells converge to an embryonic stem cell (ESC)-like state of DNAm^37^. The effect of passage on DNAm has also been examined in the context of cellular senescence, where methylation patterns in high passage cells were compared with immortalized cell lines^38,39^. In immortalized lines, DNAm undergoes stochastic changes following immortalization, whereas cells passaged to senescence underwent a programmed set of changes in DNAm consistent across samples^38,39^.

Furthermore, the effects of long-term culturing have also been explored previously in mouse intestinal IEO^40^. Specifically, Tao *et al*. found that IEO were epigenetically unstable over long term culturing. As a result, increased hypermethylation of Wnt-negative regulators led to activation of Wnt signaling rendering IEO to a ‘stem-cell’ like state and hence more permissive to tumorigenesis. Interestingly, in our study we also observed an enrichment for genes involved in Wnt signaling that were associated with CpGs found to gain DNAm (see Supplementary Information). This suggests that passage-associated epigenetic changes may contribute to altered Wnt signaling, or vice versa that continuous stimulation of this pathway via culture medium leads to these changes/induces silencing of negative regulators of Wnt signalling.

DNA methylation changes and, in particular, the global loss of DNA methylation has been observed in various malignancies including colorectal cancer. When examining culture associated DNA methylation changes in human mucosa derived IEO we observed several similarities. These include the overlap and similar directional methylation change between passage associated DMRs with known cDMRs, as well as the significant proportion of heteroskedastic methylation changes. The latter has been linked to higher variability in DNAm of colon cancer compared to normal samples^41^. Furthermore, global loss of DNA methylation in cancer has been attributed to incomplete remethylation of CpGs during mitosis possibly as a result of the higher cell turnover^33^. In keeping with this hypothesis, CpGs that lose DNAm as part of malignant transformation are frequently found in late replicating regions of the genome. This lends further support to the hypothesis that rapid cellular turnover of human IEO *in-vitro* may contribute in part to observed epigenetic changes, some share similarities to malignant cellular transformation.

There are several possible explanations for these passage changes, all of which are speculative and further works will be required to understand the mechanism of DNAm and expression change with culture duration. Regardless of the mechanism causing the observed changes in DNAm with time spent in culture, considering passage in experimental designs is an important factor. For example, when assessing potential DNAm changes during *in-vitro* differentiation of IEO, a comparison between undifferentiated and differentiated low passage IEO did not yield any significant changes suggesting DNAm is stable upon *in-vitro* differentiation. Similarly, when making the same comparison within high passage organoids, no changes are observed. However, a comparison is made between low passage undifferentiated and high passage differentiated IEO identifies 6,041 significantly differentially methylated CpGs. This example illustrates the confounding impact of culture duration on IEO culture derived experiments and further emphasises the requirement to adjust experimental set up as well as data analyses accordingly.

Our study profiled IEO up to a passage of 16, relating to approximately 5 months in culture. However, it is highly likely that longer culture durations would enhance the observed epigenetic changes and perhaps increasingly impact on cellular function. Although we have looked exclusively at IEO, it is also likely that the observed culture duration effects are not limited to IEO and would be seen in organoids derived from other tissues. Further studies are required to address these key questions arising from our work, but it is clear culture duration should be considered whenever using organoids.

Our study has identified distinct, culture associated DNA methylation changes in human mucosa derived IEO that impact gene transcription and cellular function and share features of malignant transformation. Although global epithelial cell function was found to be retained, our study highlights the critical importance of considering culture duration in the experimental design and interpretation of data derived from human IEO.

## METHODS

### Patient Recruitment and Sample Collection

Intestinal biopsies were collected from the terminal ileum (TI) and sigmoid colon (SC) from 52 children aged 1 to 16 years undergoing diagnostic endoscopy. This study was conducted with informed patient and/or carer consent as appropriate, and with full ethical approval (REC-12/EE/0482)

### Human Intestinal Epithelial Organoid Culture

Human IEO were generated from mucosal biopsies by isolation of intestinal crypts and culturing in Matrigel® (Corning) using media reported in **Table S1** as described previously^7,36,42^. The medium was replaced every 48–72 h and once the IEO were well established, they were passaged every 7-10 days by mechanical disruption and re-seeded in fresh Matrigel. IEO cultured up to passage number 4 are considered as low passaged organoids, while IEO cultured from 5 to passage number 16 are considered as high passaged organoids.

### *In-vitro* Differentiation and Co-culture with Pro-inflammatory Cytokines of Human IEO

*In-vitro* differentiation of human IEO was performed by culturing IEO in standard growth medium for four days followed by removal of Wnt agonists (referred as differentiation medium, **Table S2)** for an additional four days. For the treatment of human IEO with pro-inflammatory cytokines, IEO were cultured for 5 days after splitting in the growth medium, followed by 24 h treatment with recombinant human protein TNFα (H8916, Sigma Aldrich) at 40 ng/ml or IFNγ (PHC4031, Life Technologies) at 20 ng/ml. Bright-field images were taken using an EVOS FL system (Life Technologies).

### Human IEO Growth and Barrier Integrity Assessment

Human IEO growth was assessed using the Incucyte SX5 by imaging every 6 hours over 7 days for each passage. After 7 days, the images were analysed using the Incucyte IEO analysis software, which allowed measurement of the IEO area over time. For each human IEO line and for each passage, 6 wells were imaged and analysed to generate an average measurement of IEO growth. Comparisons between IEO passages were made using ANOVA.

Human IEO barrier integrity was evaluated culturing the IEO at early and late passage, as described above from day 0 to day 4. On day 5 IEO were collected from 48-well plates and transferred to 96-well plates, seeding 5-10 IEO per well in 5μl of Matrigel and 100μl of growth medium. Using 3 wells per condition, human IEO were cultured or in standard condition medium, or in vehicle control medium (+DMSO), or in Forskolin (5μM) medium in the presence or absence of IFNγ (20 ng/ml) or TNFα (40 ng/ml). The plates were placed in the Incucyte SX5 to be imaged every 2 hours for 48 hours. After 2 days, the experiment was stopped and images analysed to measure IEO area over time using the Incucyte IEO analysis software.

### Harvesting of Human IEO, DNA and RNA Extraction

At the end of each experiment, human IEO were harvested and both DNA and RNA were extracted using the AllPrep DNA/RNA mini kit (Qiagen, Hilden, Germany). DNA was bisulfite-converted using EZ DNA methylation Gold™ kit (Zymo Research, Irvine, CA, USA).

### DNAm Profiling and RNA Sequencing

Genome-wide DNAm was profiled using the Illumina EPIC platform (Illumina, Cambridge, UK), and deposited in ArrayExpress under accession Numbers: E-MTAB-9748 and E-MTAB-11545. An overview of sample numbers can be found in **Table 1**.

Expression profiling was performed using RNA-sequencing (RNA-seq) by Cambridge Genomic Services (University of Cambridge) and can be found in ArrayExpress under accession Number: E-MTAB-11548. Code for analysis is available at: redgar598.github.io/DNAm_organoid_passage.

### Access to Data

All authors had access to the study data and had reviewed and approved the final manuscript.

### DNAm Data Preprocessing and Quality Control

DNAm data was processed using the minfi package^43^, specifically the “preprocess” function to extract beta values from IDAT files. Data was then normalised based on control probes on each array using functional normalization^44^. Removal of two samples as outliers and those failing basic sense checks resulted in 80 IEO samples derived from 46 individuals. Starting with the 866,238 probes on the EPIC, probes were filtered if: they assay a polymorphic CpG^45^, are on a sex chromosome, had a demonstrate potential to cross hybridized to several regions of the genome^45^, or had a detection p value >0.05 in 1% of samples. This filtering left 798,096 CpGs for analysis.

### Correlation of IEO DNAm with passage

The association between DNAm and culture duration (quantified by passage) was investigated with Principal Component Analysis (PCA). The loadings of each PC were associated with technical and biological variables using ANOVA for categorical variables or Spearman correlations for continuous variables. An association between DNAm and passage was also tested on an individual CpG level. Details of computational analyses performed are provided in the supplementary methods section (Supplementary Methods). In brief, differential DNAm with increasing passage number was tested using linear models at each of 798,096 CpGs and significant heteroskedasticity in DNAm was tested with a Breusch–Pagan test.

### Public DNAm Data

Publicly available datasets used in this study are summarised in Table 1. Details of computational approaches are provided as part of the supplementary methods section (Supplement Methods)^46–48^.

### Matched DNAm and Gene Expression Cohort

The DNAm data was measured and processed as for Cohort 1 (Table 1). In the 18 untreated, undifferentiated IEO, differential DNAm with increasing passage number was tested with a linear model with a covariate for donor. For differential DNAm with differentiation and pro-inflammatory cytokine treatments, samples were split into low or high passage then within those groups a linear model with a covariate for donor was used to identify any differential CpGs.

### Enrichment DNAm Passage Changes in Genomic Features

Enrichment of CpGs that displayed DNAm changes with passage in various genomic features was tested. Analyses were performed separately for CpGs showing hyper- and hypomethylation with increased passage as well as a heteroskedastic DNAm pattern. The differential and heteroskedastic CpGs identified were explored for enrichment in genomic regulatory features. The Ensembl Regulatory Build^32^ was collected for GRCh37 using BioMart^49^ (retrieved November 2019). CpGs on the EPIC array were annotated as overlapping any of the 6 regulatory regions or as not in any annotated regulatory region. Enrichment p values for differential and heteroskedastic CpGs, in each regulatory region, were calculated using 1,000 randomly sampled lists of CpGs, to account for the underlying distribution of CpGs on the EPIC array (**Figure S2**). For hypomethylated and hypermethylated CpGs a change in DNAm of 0.15 of -0.15 was required. Therefore, the background of CpGs was modified to exclude those with a DNAm value >0.15 for hypomethylated CpGs and <0.85 for hypermethylated CpGs in the passage one IEO. These 223,695 and 295,469 CpGs, respectively, could never pass the threshold of change in DNAm and should not be included in the background CpGs list.

Similarly, enrichment p values were generated for CpGs in previously described cancer differentially methylated regions (c-DMR)^20^. Then to assess distance from origins of replication, the absolute minimum distance of a CpG from a boundary of an origin of replication (ORC2) binding peak^35^ was used. Finally, the mean distance of CpGs associated with passage was compared to the means of 1,000 randomly sampled lists of CpGs on the EPIC array, as above for regulatory region associations.

### RNA Sequencing Data Analyses

For each of the 42 validation samples, RNA was prepared with the Truseq mRNA library preparation (Illumina, San Diego, CA, USA) and sequencing was performed on NextSeq 75 cycle high output. RNA-seq data was quality controlled using FastQC^50^. Reads were pseudoaligned using kallisto^51^ indexed human transcriptome (GRch38) and quantified with 100 bootstraps. Using sleuth^52^ differential expression was measured at the gene level by aggregating across all transcripts associated with a gene (Ensembl Genes 104)^53,54^. Gene expression was associated with passage as a continuous measure (2-12 passages) using a likelihood-ratio-test with a covariate for donor. For differential expression with differentiation and pro-inflammatory cytokine treatments, samples were split into low or high passage then within those groups a likelihood-ratio-test with a covariate for donor was used with an FDR < 0.05 considered significant.

## Acknowledgements

Work described in this publication was supported by grants from the Medical Research Council (MRC, New Investigator Research Grant), European Society of Paediatric Gastroenterology, Hepatology and Nutrition (ESPGHAN Networking grant), Guts UK charity and British Society for Paediatric Gastroenterology, Hepatology and Nutrition (BSPGHAN). SL was supported by the UCLA Broad Stem Cell Research Center Predoctoral Training Grant and the Rose Hills Foundation. DLJ was supported by the UCLA Broad Stem Cell Research Center Innovation Award and the Rose Hills Foundation, the CURE: Digestive Diseases Research Center at UCLA Pilot and Feasibility Study Award (center grant P30 DK 41301), and the UCLA CTSI/BSCRC/DGSOM Regenerative Medicine Theme Award. JK was supported by Crohn’s in Childhood Research Organization (CICRA). DRZ and RDE were supported by core funding from the European Molecular Biology Laboratory (EMBL).

## Author Contributions

RDE, MZ and DRZ designed the study, formulated the research questions and wrote the manuscript. RDE performed all data analyses supported by DRZ. FP, KMN, ARF and RH contributed to data curation and experimental design. KMN, JK, FP, ARF and AC performed experimental work. FT, CS, RH and KOH recruited patients, collected biopsies and contributed to documentation of clinical information. DLJ and SL provided data and clinical information for an independent validation cohort and contributed to critical discussion of the findings and editing of the final manuscript. All authors read and approved the final manuscript. MZ and DRZ contributed equally to this work.

