## Supplementary Information for "Culture Associated DNA Methylation Changes Impact on Cellular Function of Human Intestinal Organoids"

April 1, 2022

**Table S1:** List of growth medium components

| **Basal Organoid medium = ADF+++<br>Products from Gibco by Thermo Fisher Scientific** | **Final concentration** |
|---|---|
| Advanced DMEM/F12 (ADF) | 1x |
| GlutaMax | 2 mM |
| HEPES buffer | 10 mM |
| Penicillin/Streptomycin | 0.5 U/ml |
| **Complete medium (total volume 50ml):** | **Final concentration** |
| ADF+++ (see above) | 76,3% (vol/vol) |
| Wnt surrogate-Fc Fusion Protein (ImmunoPrecise) | 0,2 nM |
| R-spondin-1 conditioned medium | 20 % (vol/vol) |
| Primocin (Invivogen San Diego, CA, USA) | 500 $\mu$g/mL |
| B-27® Supplement (Invitrogen, Carlsbad, CA, USA) | 1x |
| Nicotinamide (Sigma, St. Louis, MO, USA) | 10 mM |
| N-Acetylcysteine (Sigma, St. Louis, MO, USA) | 1.25 mM |
| A3801 (Tocris, Bristol, UK) | 500 nM |
| SB202190 (Sigma, St. Louis, MO, USA) | 10 $\mu$M |
| Murine EGF (Invitrogen, Carlsbad, CA, USA) | 50 ng/mL |
| Murine Noggin (Peprotech, Rocky Hill, NJ, USA) | 100 ng/mL |

**Table S2:** List of differentiation medium components

| **Differentiation medium (total volume 12.5ml):** | **Final concentration** |
|---|---:|
| ADF+++ (see above) | 97.44% (vol/vol) |
| Primocin (Invivogen San Diego, CA, USA) | 500 $\mu$g/mL |
| B-27® Supplement (Invitrogen, Carlsbad, CA, USA) | 1x |
| N-Acetylcysteine (Sigma, St. Louis, MO, USA) | 1.25 mM |
| A3801 (Tocris, Bristol, UK) | 500 nM |
| SB202190 (Sigma, St. Louis, MO, USA) | 10 $\mu$M |
| Murine EGF (Invitrogen, Carlsbad, CA, USA) | 50 ng/mL |
| Murine Noggin (Peprotech, Rocky Hill, NJ, USA) | 100 ng/mL |

**Table S3. Top over-represented GO groups in the list of genes adjacent to heteroskedastic CpGs.**Columns are: GO ID, name of the GO gene set, number of genes annotated to that gene set and associated to a heteroskedastic CpG, Benjamini-Hochberg corrected enrichment p value (which the data is sorted by), Multifunctionality (MF) score and the names of genes annotated to that gene set and associated to a heteroskedastic CpG.

**Table S4. Top over-represented GO groups in the list of genes adjacent to hypomethylated CpGs.** Columns are: GO ID, name of the GO gene set, number of genes annotated to that gene set and associated to a hypomethylated CpG, Benjamini-Hochberg corrected enrichment p value (which the data is sorted by), Multifunctionality (MF) score and the names of genes annotated to that gene set and associated to a hypomethylated CpG.

**Table S5. Top over-represented GO groups in the list of genes adjacent to hypermethylated CpGs.** Columns are: GO ID, name of the GO gene set, number of genes annotated to that gene set and associated to a hypermethylated CpG, Benjamini-Hochberg corrected enrichment p value (which the data is sorted by), Multifunctionality (MF) score and the names of genes annotated to that gene set and associated to a hypermethylated CpG.

**Table S6. Genes differentially expressed and DNAm with passage.** Columns are: gene name, Ensembl gene ID, p value and q value for gene expression association to passage, CpGs associated to the gene which were hypermethylated or hypomethylated.

**Table S7. Genes differentially DNAm with passage and differentially expressed with differentiation but dependent on passage group.** Columns are: gene name, Ensembl gene ID, low passage p value and q value for gene expression association to differentiation, high passage p value and q value for gene expression association to differentiation, CpGs associated to the gene which were hypermethylated or hypomethylated.

**Table S8. Genes differentially DNAm with passage and differentially expressed with IFN$\gamma$ treatment but dependent on passage group.** Columns are: gene name, Ensembl gene ID, low passage p value and q value for gene expression association to IFN$\gamma$ treatment, high passage p value and q value for gene expression association to IFN$\gamma$ treatment, CpGs associated to the gene which were hypermethylated or hypomethylated.

**Table S9. Genes differentially DNAm with passage and differentially expressed with TNF$\alpha$ treatment but dependent on passage group.** Columns are: gene name, Ensembl gene ID, low passage p value and q value for gene expression association to TNF$\alpha$ treatment, high passage p value and q value for gene expression association to TNF$\alpha$ treatment, CpGs associated to the gene which were hypermethylated or hypomethylated.

**Video File 1.** Bright-field videos of TI derived IEOs at low and high passage, cultured in Forskolin, IFN$\gamma$+Forskolin or TNF$\alpha$+Forskolin medium, taken by Incucyte; scale bars= 900$\mu$m.

### Supplementary Methods

#### Correlation of IEO DNAm with passage

Each of the 798,096 CpGs were tested for both heteroskedasticity in DNAm and differential DNAm with increasing passage number. As there are more samples of organoids with one to four passages (Figure 1E), low passage samples (passages 1-4) were downsampled to five samples per passage level. The downsampling was done 1,000 times, and each cohort subsample contained 42 organoid samples for testing. A linear model was fit to measure the association between DNAm and passage in each subsample, at each CpG. In addition to the differential p value, the change in DNAm between organoid samples from 1 and 16 passages was calculated as a delta beta. In each cohort subsample a Breusch–Pagan test on the residuals of the linear model between passage and DNAm was used to test for heteroskedasticity. The number of times, out of 1,000, the differential or heteroskedasticity p values were significant (FDR $< 0.05$) was used to generate a robust differential and heteroskedasticity p value for each CpG. Heteroskedastic CpGs were defined as those passing the Breusch–Pagan test in a significant number of the 1,000 subsamples ($p<0.05$). Similarly, differentially DNAm CpGs were defined as those with an association between passage and DNAm in a significant number of subsamples ($p< 0.05$) but also with an absolute delta beta greater than 0.15. All samples were submitted to the epigenetic clock software[28] and predicted epigenetic age was compared to chronological age of the patient when the biopsy used to derive the IEO was taken.

#### Public DNAm Data

To validate the passage effect, more intestinal epithelial IEO were collected from publicly available data (Table 1). The data deposited under E-MTAB-4957[19] was collected as IDAT files from ArrayExpress and we refer to this as Cohort 2. The data were processed in the same way described above, except with a different annotation as Cohort 2 is on the 450K array not the EPIC [29]. Of the 42 available pediatric IEO, again from the SC and TI, 30 are unique IEOs samples but 12 IEOs were derived from individuals also used in our cohort. These 12 IEOs were excluded. To associate DNAm at individual CpGs to passage, a linear model was used. However, in the Cohort 2 IEOs the downsampling of the early passage IEOs was not done since there are fewer IEOs available, making downsampling challenging. Nonetheless, the available IEOs do have a more even spread of samples across passages.

Another dataset of IEO is available under GSE141256[10], these were collected as IDAT files from Gene Expression Omnibus (GEO) [30]. We refer to this as Cohort 3. In addition, through a personal communication, passage number for these samples was obtained. The DNAm data were processed in the same way described above for both the EPIC and 450K arrays as Cohort 3 includes both. Only the 384,188 CpGs on both arrays, after probe filtering, were used. Batch correction for array type was done using ComBat[31]. In the 21 available IEO, from duodenum, jejunum or colon, linear models were used to associate DNAm to passage as in the Cohort 1 IEO.

#### Pathways Affected by DNAm Passage Changes

To explore possible pathways systematically affected by passage, the CpGs differentially DNAm with passage were associated with genes, and these genes were then tested for enrichment in gene ontology (GO) gene sets. The heteroskedastic and differential CpGs were associated with adjacent genes. A CpG was assigned to a gene based on proximity to a transcript from Ensembl Genes 99 GRCh37.p13 collected from BioMart[33]. CpGs were associated with a gene if they were located between 1500bp upstream of the transcript start and 300bp downstream of the transcript end. CpG to transcript associations were then aggregated by Gene stable ID. Then these CpG to gene associations were used for enrichment of gene ontology (GO) terms. GO annotations of the 25,676 genes associated with the EPIC probes were used as the background list. Enrichment of GO terms in the list of passage associated genes (Heteoskedastic: 13,267 genes, Hypomethylated: 5,579 genes, Hypermethylated: 3,673 genes) were tested using overrepresentation analysis in ErmineJ[55]. Significance of a GO term is reported as false discovery rates (FDR) computed using the Benjamini–Hochberg method in ErmineJ. Also included are the multifunctionality scores of GO terms[56].

Hypomethylated and heteroskedastic CpGs were enriched in similar GO gene sets. Of the 363 and 242 significantly enriched GO gene sets (FDR $< 0.05$) in heteroskedastic and hypomethylated CpGs, respectively, 147 were overlapping (Table S3 and S5). These include gene sets for adherens junction organization (GO:0034332) and cell-cell adhesion via plasma-membrane adhesion molecules (GO:0098742) or cell-cell adhesion mediated by cadherin (GO:0044331), which together could suggest consequences for IEO barrier function. However there were many unexpected GO sets as well, such as neuron projection guidance (GO:0097485) and synapse assembly (GO:0007416). Enrichment of these gene sets could be because the genome is globally hypomethylated and the genes enriched are not specific to epithelial functional pathways. Alternatively, there may be unknown functions for these genes in gut epithelial cells.

The results for hypermethylated CpGs were more meaningful. Enriched GO terms include: columnar/cuboidal epithelial cell differentiation (GO:0002065) and glandular epithelial cell differentiation (GO:0002067) (Table S5). Interestingly several Wnt signalling gene sets are enriched for hypermethylated genes (e.g. GO:0030111 and GO:0007223)

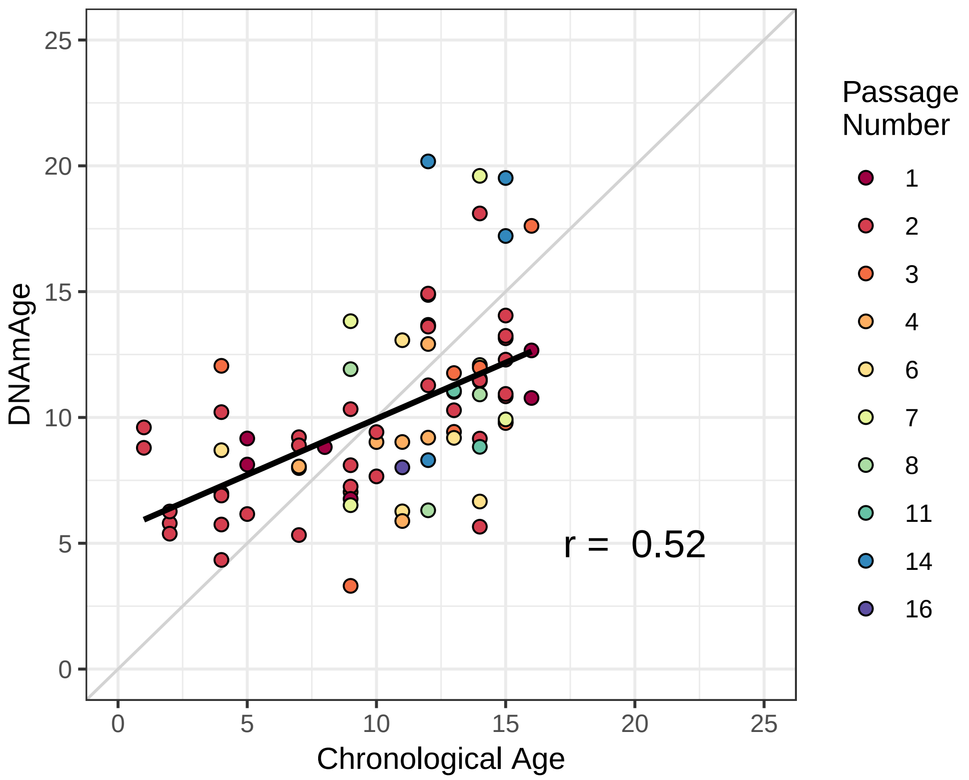


**Figure S1: Chronological age and epigenetic age are similar in patient derived IEO.** The association between chronological age and epigenetic age is shown, with points coloured by the passage of the IEO.

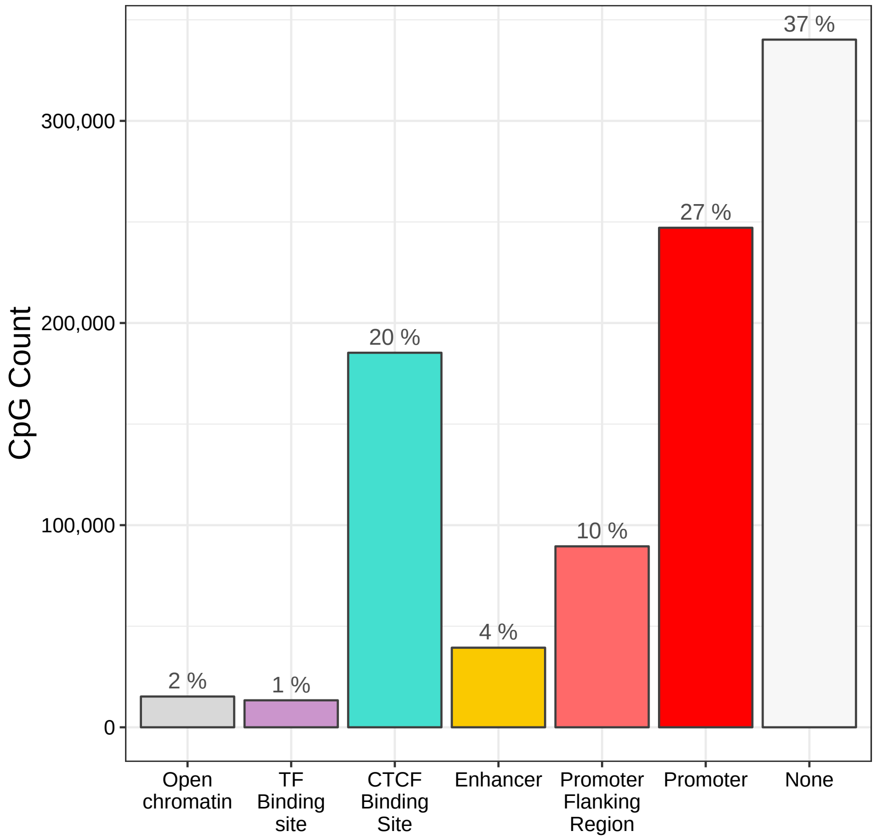


**Figure S2: Background distribution of CpGs on the EPIC array.** Number of 800,383 EPIC array CpGs (used in this analysis) in regulatory genomic features.

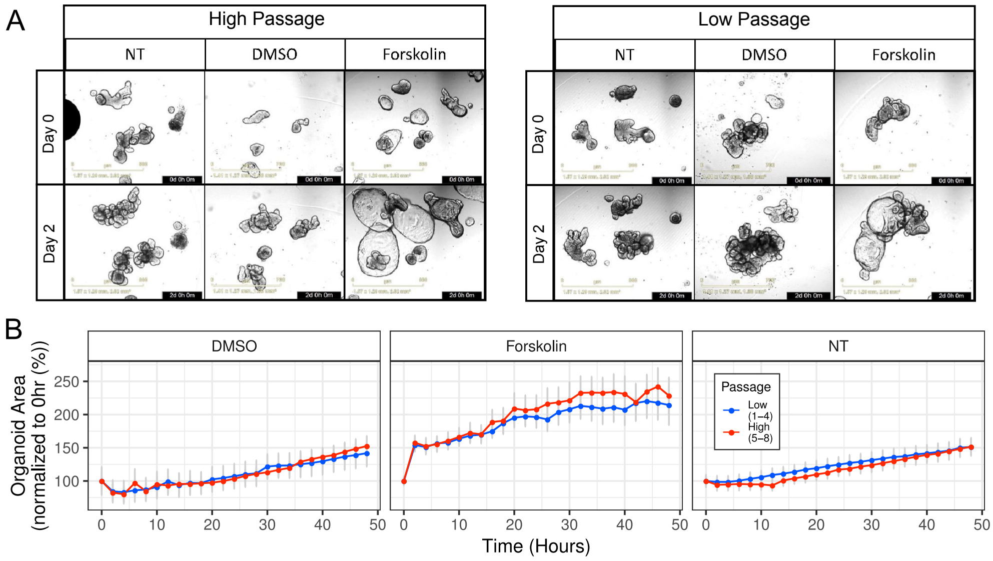


**Figure S3: High and low passage IEO look respond similar forskolin plus cytokine treatments.** (A)Representative bright-field images of TI derived IEOs at high and low passage, cultured in standard medium (NT), vehicle control medium (DMSO) and Forskolin, taken by Incucyte at two different time points (day0 and day2); scale bars= 800$\mu$m. (B) Comparison of the organoids area between early and late passage organoids cultured in DMSO, Forskolin and standard medium (n=2, 4 technical replicates per each biological replicate).

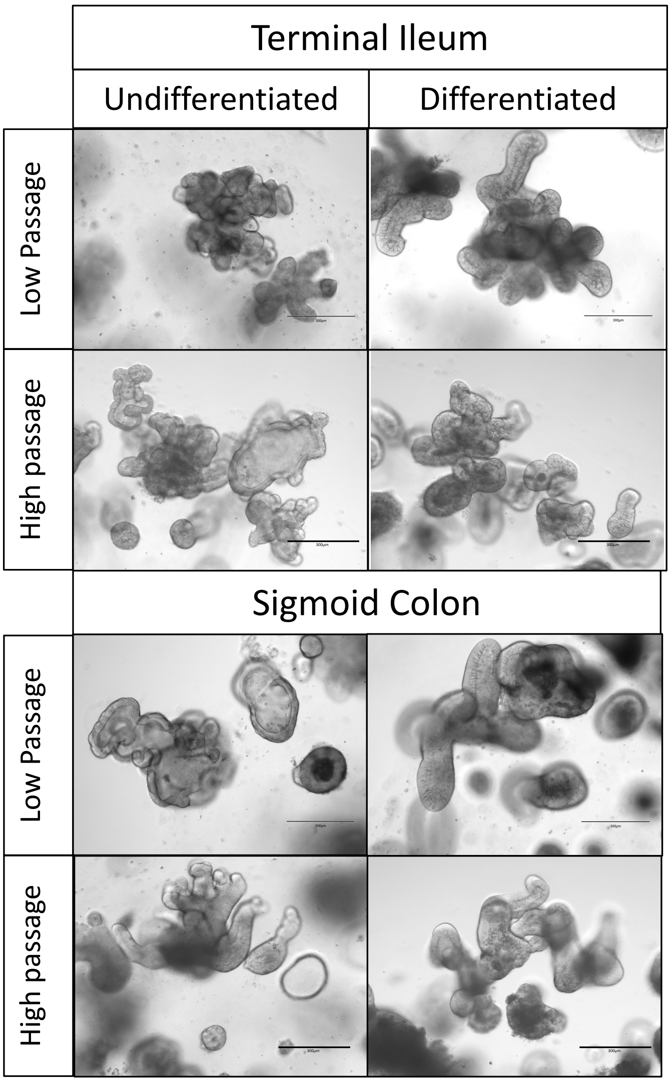


**Figure S4: High and low passage IEO look similar before and after differentiation.** Bright-field images of TI and SC derived IEOs following *in vitro* differentiation respectively, taken by EVOS FL system (Life Technologies), scale bars= 300$\mu$m.

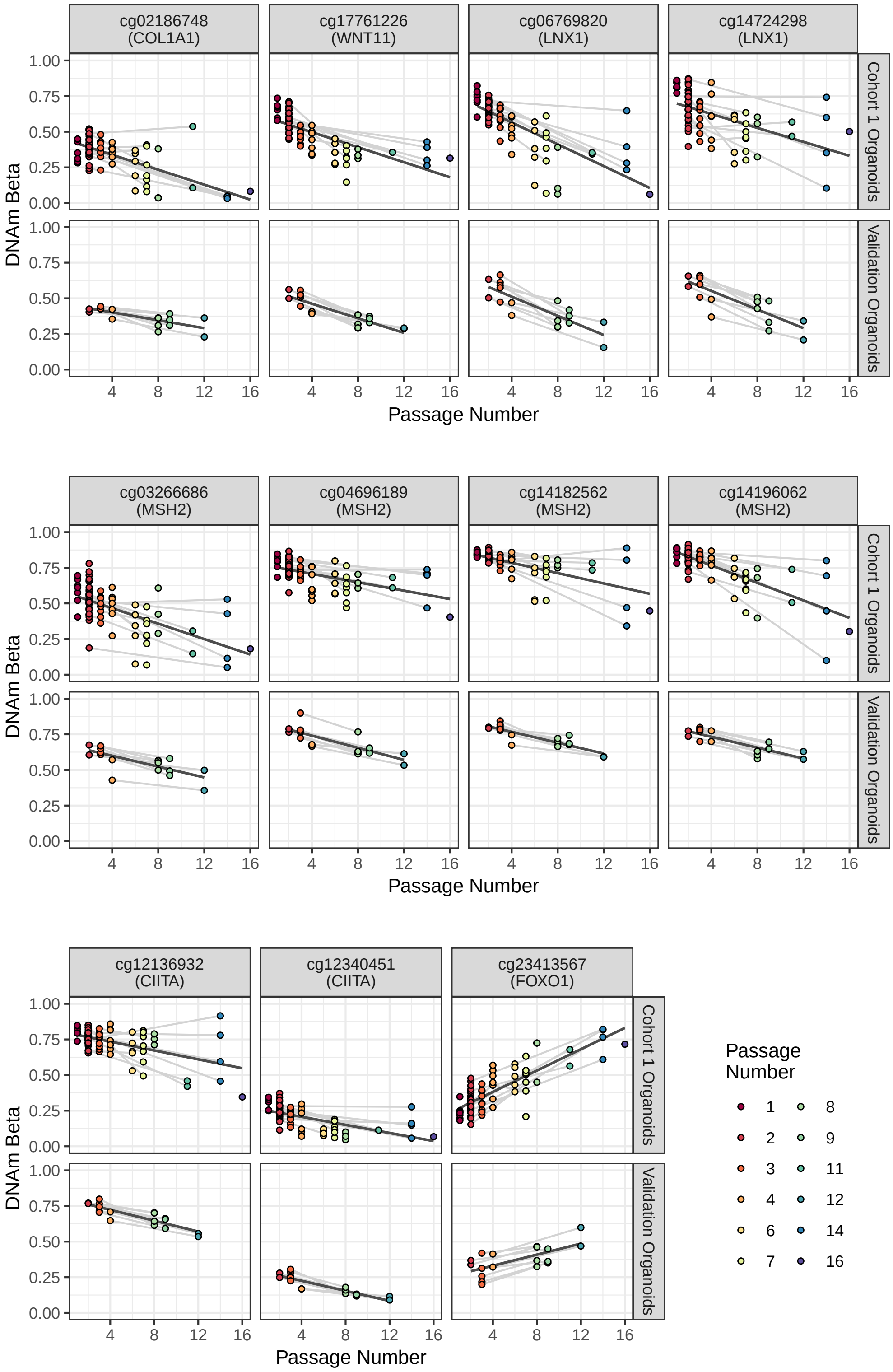


**Figure S5: Differential DNAm with passage at CpGs associated to genes with expression dependent on IEO passage.** CpGs with DNAm significantly associated to passage in Cohort 1 and validation IEO. Samples are coloured by passage number and grey lines connect samples derived from the same patient. Regression lines between passage and DNAm are in black.
