## Supplementary figures and images for "Culture Associated DNA Methylation Changes Impact on Cellular Function of Human Intestinal Organoids"

### Supplementary Video 1

#### Slide 1
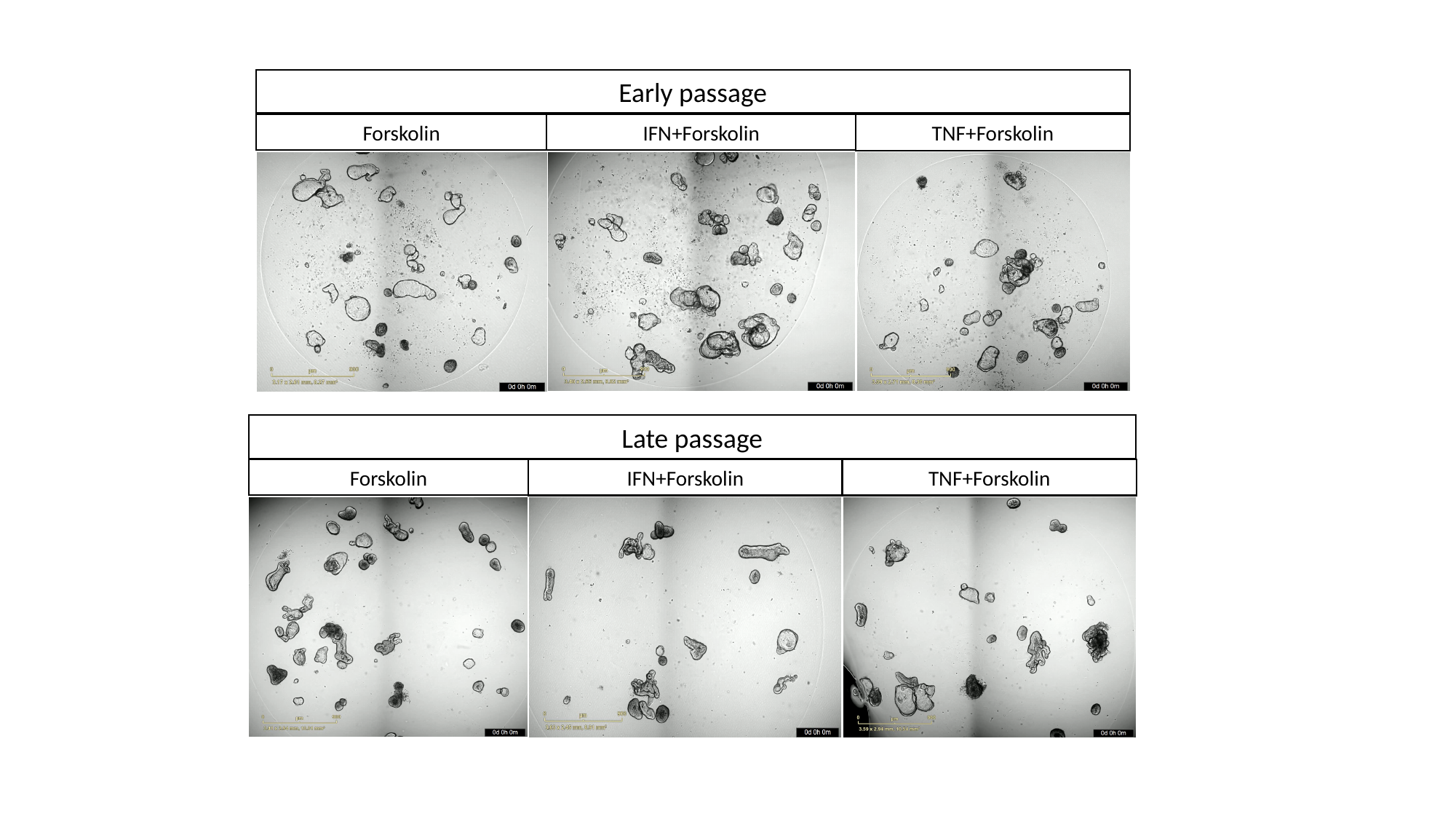

Early passage
Forskolin
Late passage
Forskolin
